# A transcriptome atlas of zygotic and somatic embryogenesis in Norway spruce

**DOI:** 10.1101/2024.04.13.589382

**Authors:** Katja Stojkovič, Camilla Canovi, Kim-Cuong Le, Nicolas Delhomme, Ulrika Egertsdotter, Nathaniel R. Street

## Abstract

Somatic embryogenesis (SE) is a powerful model system for studying embryo development and an important method for scaling up availability of elite and climate-adapted genetic material of Norway spruce (*Picea abies* L. Karst). However, there are several steps during the development of the somatic embryo (Sem) that are suboptimal compared to zygotic embryo (Zem) development. These differences are poorly understood and result in substantial yield losses during plant production, which limits cost-effective large-scale production of SE plants. This study presents a comprehensive data resource profiling gene expression during zygotic and somatic embryo development to support studies aiming to advance understanding of gene regulatory programmes controlling embryo development. Transcriptome expression patterns were analysed during zygotic embryogenesis (ZE) in Norway spruce, including separated samples of the female gametophytes and Zem and at multiple stages during SE. Expression data from eight developmental stages of SE, starting with pro-embryogenic masses (PEMs) up until germination, revealed extensive modulation of the transcriptome between the early and mid-stage maturing embryos and at the transition of desiccated embryos to germination. Comparative analysis of ZE and SE identified differences in timing relative to SE development and functional enrichment of these provided insight into the associated biological processes. Orthologs of transcription factors known to regulate embryo development in angiosperms were differentially expressed during Zem and Sem development and in the different zygotic embryo tissues, providing clues to the differences in development observed between Zem and Sem. This resource represents the most comprehensive dataset available for exploring embryo development in conifers.

**Significance Statement:** Somatic embryogenesis is used as a model system to study embryo development, however detailed information to verify similarities and explain differences between somatic and zygotic embryogenesis is largely missing for conifers. This data resource provides sequential mRNA transcriptome data from nine stages of conifer zygotic embryo and female gametophyte development, and eight stages of somatic embryo development, to enable exploration of biological questions and comparisons of the two developmental processes.

## Introduction

The gymnosperm and angiosperm lineages diverged about 300 million years ago, resulting in different morphological and functional pathways for key steps of development such as embryogenesis (La Torre et al. 2019). Characteristic elements of conifer embryo development include the occurrence in many species of a polyembryonic process in which a multitude of immature embryos develop before one embryo becomes dominant (Arnold et al. 2019). The process of polyembryony has been suggested to underlie the ability to induce the *in vitro* somatic embryogenesis (SE) process in conifers whereby multiplication of the early-stage embryos (pro-embryogenic masses; PEMs) can be prolonged for extended periods, generating unlimited numbers of somatic embryos (Sem) from the original zygotic embryo (Zem). The high similarity between zygotic embryogenesis (ZE; Cairney and Pullman 2007) and SE in conifers renders SE a powerful model system to study embryogenesis, for example to elucidate gene expression regulatory mechanisms (reviewed in Trontin et al. 2016). A major strength of SE as a model system is the ability to produce an almost unlimited number of Sem of the same genotype that can be sampled at distinct, synchronised stages of the developmental process. SE is particularly effective in *Picea* species as the process works efficiently across many genotypes (Mamun et al., 2018). This strong potential for clonal multiplication additionally makes SE a valuable tool for the forestry industry to obtain clonally propagated conifer plants derived from elite varieties where clonal propagation by cuttings or other techniques is limited or absent (Egertsdotter et al. 2019).

In *P. abies*, Zem development starts in the same year after pollination and continues until a fully mature, desiccation-tolerant embryo has formed eight to twelve weeks later. For most conifers, embryo development progresses through three stages: proembryo, early embryogeny and late embryogeny (Singh 1978). Proembryo formation is characterised by a free nuclear stage ending when the proembryo elongates into the corrosion cavity of the female gametophyte (FG), followed by the further elongation of the suspensor mass and development of a generative root meristem during early embryogeny. Root and shoot meristem development are concluded during late embryogeny (Singh 1978). During seed development, programmed cell death of FG cells adjacent to the embryo also provides nutrients for the growing embryo (Durzan 2012). FG tissue further supports the onset of germination by allocating storage reserves to the developing Zem and remains active until seed imbibition (Vuosku et al. 2009).

The *in vitro* SE process in conifers is typically started from an immature Zem that will produce PEMs in response to auxin and cytokinin. Recently, low frequency of Sem initiation from shoot buds of *P. abies* was however demonstrated from four- to six-year-old plants that had originated from Sem (Varis et al. 2018). When the PEMs have formed a callus-like mass that can be captured for continued multiplication, maturation can be induced by replacing the cytokinin and auxin with abscisic acid (ABA) and an increased osmotic potential. The fully mature somatic embryos are often subjected to a period of drying by exposure to a relatively drier gas phase to become partially desiccated before the onset of germination. After germination, embryos showing root and shoot development can be acclimated to *ex vitro* conditions under controlled humidity and temperature (Egertsdotter 2018).

SE plants are attractive for use in forestry applications as seeds from elite crossings can be clonally propagated and directly provide elite planting material for deployment (clonal propagation of families) or utilized for testing in breeding programs and for subsequent deployment for clonal forestry (Rosvall et al. 2019). The SE process in conifers is slow and there can be loss of genotypes in addition to limitations in the number of embryos produced at each step, most notably at the stages of initiation of PEMs from the Zem, limitations to multiplication rates and conversions between PEMs to mature embryos, the transition of mature embryo to germinants and, finally, plant growth and establishment from the germinant. Efforts to optimize SE method protocols have rendered some improvements but further optimizations are required to enable cost-effective SE plant production across genotypes. In particular, the formation of a fully mature, desiccation tolerant embryo is still a limiting factor in most conifer species. Compared to the Zem, the Sem lacks the FG supporting tissue and, arguably, components of the FG are missing from the SE culture medium. The composition and metabolism of the FG tissue have also been analysed to improve the SE processes by amending the *in vitro* culture medium and methods to resemble the FG environment of the Zem more closely (reviewed by Pullman and Buccalo 2014).

While gene expression during embryo development has been extensively studied in angiosperm model systems, efforts to analyse global transcript abundance during conifer embryo development have been hampered by a lack of reference genome assemblies due to the large genome sizes of gymnosperms, which average 200 times the size of the Arabidopsis genome (Mackay et al. 2012). However, it has been demonstrated that gene families active during embryo development in angiosperms are also active in conifers, with differential expression (DE) of epigenetic regulators and transcription factors (TFs) implicated as developmental regulators in angiosperms being detected by microarray analysis in Zem of *Pinus pinaster* (de Vega-Bartol et al. 2013), and RNA sequencing of Zem and FG respectively of *Pinus sylvestris* (Merino et al. 2016) and whole seeds of *Picea mongolica* (Yan et al. 2021). Efforts to analyse global transcript abundance during conifer embryo development have otherwise mainly focused on SE material due to the easy access of embryos at different developmental stages and the possibilities to analyse embryos under different growth conditions and from cell lines with varying degrees of developmental arrest (reviewed in Trontin et al. 2016). SE cultures composed of early-stage embryos (PEMs) also offer the advantage of being amenable to genetic transformation to test the function of candidate genes with regulatory functions during embryo development (e.g. Vetrici et al. 2021) and have recently been shown to be amenable to genomic modifications via CRISPR-Cas, permitting more in-depth analyses of developmental regulators of embryo development in *Pinus radiata* (Poovaiah et al. 2021) and *Picea glauca* (Cui et al. 2021).

Transcript profiling of conifer zygotic and somatic embryos at corresponding developmental stages has also been conducted with the aim of identifying discrepancies between the respective developmental stages. The SE process for scalable plant production in conifers is suboptimal in most, and blocked in some, species. As such, learning from the natural processes during ZE could help identify approaches to improve SE plant production methods. This approach was explored early on by using a 326 cDNA fragment macroarray in *Pinus taeda* to compare the most mature stage of Sem to the full series of developmental stages of Zem. From this comparison of a limited number of gene fragments it appeared that the most mature stage reached by the Sem *in vitro* did not correspond to the same level of maturity as Zem, suggesting that Sem development in *P. taeda* was blocked at a premature stage (Cairney et al. 2000, Pullman et al. 2003). More recently, gene expression in Zem and FG at three different developmental stages during seed development was compared in two cell lines with different Sem maturation competency in *Araucaria angustifolia* (Elbl et al. 2015). The results from the comparison of gene expression indicated that the cell line with blocked embryo maturation had a disturbed auxin distribution pattern.

In the present study, a comprehensive developmental series of *P. abies* Zem and their associated FG were profiled by RNA-Sequencing (RNA-Seq) at biweekly intervals. Nine time points were covered, starting two weeks after pollination, and ending with a fully mature and desiccated embryo. These data cover more developmental stages than any previous study and provide new information of gene expression events during the very first stages of seed development in addition to key events such as desiccation. Furthermore, eight developmental stages of somatic embryos were profiled by RNA-Seq and tentatively aligned with corresponding stages of zygotic embryos based on embryo morphologies and differential gene expression results. Extensive modulation of the transcriptome at specific points during SE and ZE development was detected, with DE analysis conducted to identify candidate genes and biological processes involved in the *P. abies* embryo development transcriptional program. There was extensive DE of genes at the major developmental stage transitions, with the largest changes occurring between the early and the mid stage maturing somatic embryos and at the transition of desiccated embryos to germination. Overall, major differences in gene expression patterns were detected between Zem and Sem development, revealing differences in the two developmental processes. Furthermore, differences in expression patterns of TFs with key functions in angiosperms were detected during Zem and Sem development.

The data presented here have been integrated into PlantGenIE.org (Sundell et al. 2015) to serve as a resource for fundamental studies into the mechanisms of conifer embryo development. The resource represents a reference gene expression atlas for conifer embryo development, enabling exploration of gymnosperm embryo, understanding of which is currently lagging behind that of angiosperm species. Furthermore, the resource enables community utilisation to explore discrepancies between Sem and Zem development to improve understanding of observed discrepancies between somatic and zygotic embryogenesis. Such knowledge should facilitate development of improved methods to obtain high yields of SE plants for research and reforestation purposes.

## Results and Discussion

### Overview of gene expression during ZE

RNA sequencing at nine stages of ZE, from the seed two weeks after pollination to a fully desiccated seed, was performed to assay the transcriptome during ZE (Figure 1A). Previous work in *Picea engelmannii* × *glauca* following development of the FG and the post fertilisation events described by ultrastructural analyses revealed that fertilisation takes place around two weeks after pollination (Owens and Molder 1984). A similar time frame was shown for seed development in *P. abies* (Hakman 1993), suggesting that the first developmental stages of seeds sampled in this study two weeks after pollination would correspond to seeds around the time when fertilisation occurred, followed by proembryo development. The second sample would then contain proembryos represented by the free-nuclear stage of the zygote. Proembryogeny lasts for about a week in both *P. glauca* and *P. engelmannii* (Owens and Molder 1984) but has been observed to last for around two weeks in *P. abies* (Hakman 1993) and ends when the embryonal cells are pushed into the corrosion cavity by the suspensor cells to start early embryogeny (Singh 1978). The appearances of the female gametophytes (FGs) of the seed samples were strikingly different between the two first developmental stages and the third stage, which was considerably larger and had changed from a transparent appearance to opaque, whitish colouring. This supports the assumption that for this present study the first two stages corresponded to proembryogeny and that by the third collection time, seeds had started to transition to early embryogeny. Further support comes from the observation of the Zem resembling the typical early embryo stages with a distinctive embryo -proper at the fourth collection time (Figure 1A). The phase of early embryogeny then continues until late embryogeny starts about eight weeks later (Owens and Molder 1979). The appearances of the sampled FGs remained the same from the third stage to the last stage, which contained the fully mature and desiccated embryo. Examination of a principal component analysis (PCA) plot revealed that samples from FG and Zem clearly separated by tissue type and showed a time dependent pattern until stage Z6 (Figure 1B). Samples from the latest stages (Z7-Z9) clustered together, indicating that no major transcriptional changes occurred after stage Z7 suggesting that these stages correspond to the desiccated and dormant seed. Samples from the first two stages (Z1 and Z2) were more similar to each other than to the third stage (Z3), which clustered closer to the FG samples. All three stages (Z1-Z3) represented the whole seed before it was possible to separate FG from Zem.

**Figure 1.**
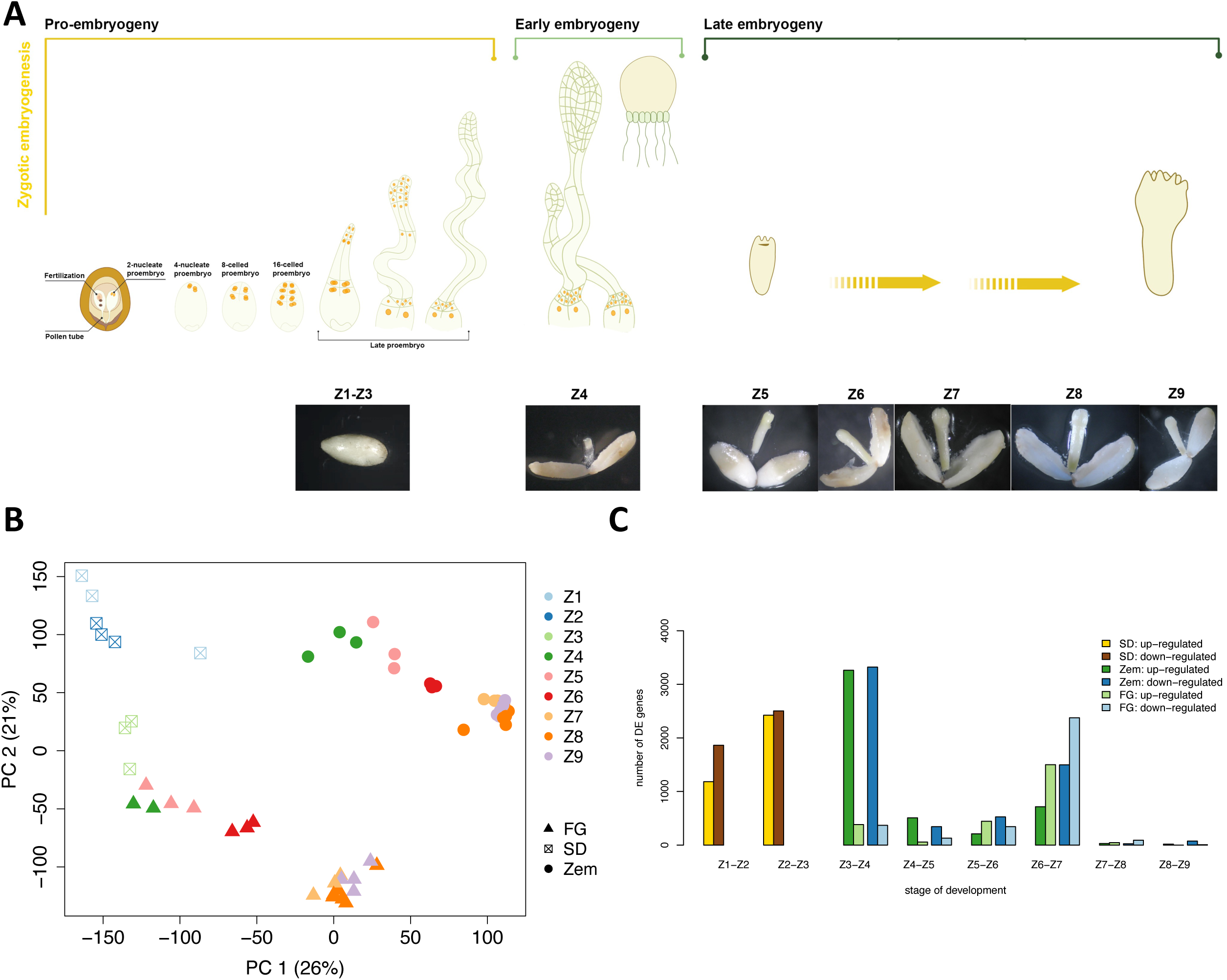
**A** Diagrammatic representation of the sampled stages during zygotic embryogenesis. Representative photos of each sampled stage are shown below the diagram. Each sample consisted of a pool of hundreds of seeds/embryos. **B** Principal component analysis plot of the RNA-Sequencing data showing the first two principal components. Different symbols indicate sample type and colours indicate sample stage. **C** Bar graph representation of the number of differentially expressed genes (FDR-adjusted p-value <0.01 and |log2 fold change| ≥0.5). Darker shades indicate down-regulated genes and lighter shades up-regulated genes. Colours indicate sample type. FG, female gametophyte; Zem, zygotic embryo; SD, seed.

Across all sampled ZE tissue types (seed, zygotic embryo and female megagametophyte) 36718 genes had detectable expression of which 40.0 % (14686) were differentially expressed (DE) in at least one stage transition in one of the tissues. Differentially expressed genes (DEGs) at any time in the seed, Zem and FG represented 19.7 % (7,230 genes), 23.8 % (8749 genes) and 13.8 % (5,050 genes), respectively, of all ZE expressed genes. The largest numbers of DEGs in ZE (Figure 1C) were observed firstly in the early stages of seed development, from Z1-Z4 (3046, 4928, 7048 DEGs between Z1-Z2, Z2-Z3, and Z3-Z4, respectively), which is when the transition from proembryogeny to early embryogeny is expected to take place (Owens and Molder 1984). In *Picea abies,* the optimal time for SE initiations from the Zem is around stage Z4, suggesting that some of the observed DEGs play a role in the dedifferentiation process when PEMs are formed. The next stage showing the largest numbers in DEGs was between Z6 and Z7 (4,940 DEGs), which is around the time when the switch from early embryogeny to late embryogeny occurs (Owens and Molder 1984). There were 792 and 212 DEGs common to the two stage transitions with the largest number of DEGs (Z3-Z4 and Z6-Z7) in Zem and FG, respectively.

The FG supports embryo development by allocating storage reserves to the developing embryo until the onset of germination and stays active until seed imbibition, as observed in *Picea glauca* (He and Kermode 2003) and *Pinus silvestris* (Vuosku et al. 2009). This is in agreement with the results from the functional enrichment analysis in the present study (discussed below). There were 5807 DEGs between Zem and FG considering all stage comparisons, with more than 1500 DEGs between the two tissue types at every time point. When comparing separated tissues of the seed from stage Z4 to the combined seed sample (Z3), a larger number of DEGs was observed for the Zem than for FG. This suggests that before Z4 the assayed transcriptome was dominated by FG and that FG shows little change in the transcriptome during these stages. In contrast, at the time of the second biggest change in the number of DEGs in the Zem (Z6-Z7), presumably corresponding to accumulation of storage compounds in preparation for desiccation, there were more DEGs in the FG. Functional enrichment analysis of the 5807 DEGs between the FG and Zem at any point of the experiment identified significant enrichment for cellular nitrogen compound metabolic process (GO:0034641, padj = 5.95e-11), multiple terms connected to chromatin organisation (GO:0006333, padj = 3.47e-8; GO:0071103, padj = 5.25e-6; GO:0051276, padj = 1.28e-4; GO:0034401, padj = 4.11e-3) and terms connected to transcription and translation (GO:0006261, padj = 3.28e-3; GO:0022613, padj = 8.79e-4; GO:0016072, padj = 2.62e-3; GO:0010467, padj = 3.64e-3).

Functional enrichments of DEGs at the major points of transition in both tissue types were also examined to gain insight into the key biological processes active (Table S1). For Zem there was significant enrichment of GO:0051276 (padj = 5.12e-7), GO:0046939 (padj = 1.77e-6), GO:0009058 (padj = 1.23e-4), GO:0016072 (padj = 2.31e-4), GO:0006323 (padj = 2.64e-4) at the Z3-Z4 transition, representing chromosome organization, nucleotide phosphorylation, biosynthetic process, rRNA metabolic process, and DNA packaging, suggesting active transcription, changes to organisation of chromatin and biosynthesis. At Z6-Z7 there was enrichment for lipid metabolic process (GO:0006629, padj = 1.15e-4), organophosphate metabolic process (GO:0044281, padj = 6.98e-4), and carbohydrate metabolic process (GO:0005975, padj = 5.474-3). In the FG, the most marked change in the transcriptome occurred at Z6-Z7, with a clear bias towards down-regulation of transcripts. These DEGs were enriched for carbohydrate metabolic process (GO:0005975, padj = 8.55e-10), cell wall organization or biogenesis (GO:0071554, padj = 1.01e-4), and external encapsulating structure organization (GO:0042431, padj = 1.07e-4).

### Overview of gene expression during SE

RNA sequencing to profile gene expression was performed at eight stages of SE in Norway spruce, from the earliest stage in culture, composed of PEMs, to an established growing plant (Figure 2A).

**Figure 2.**
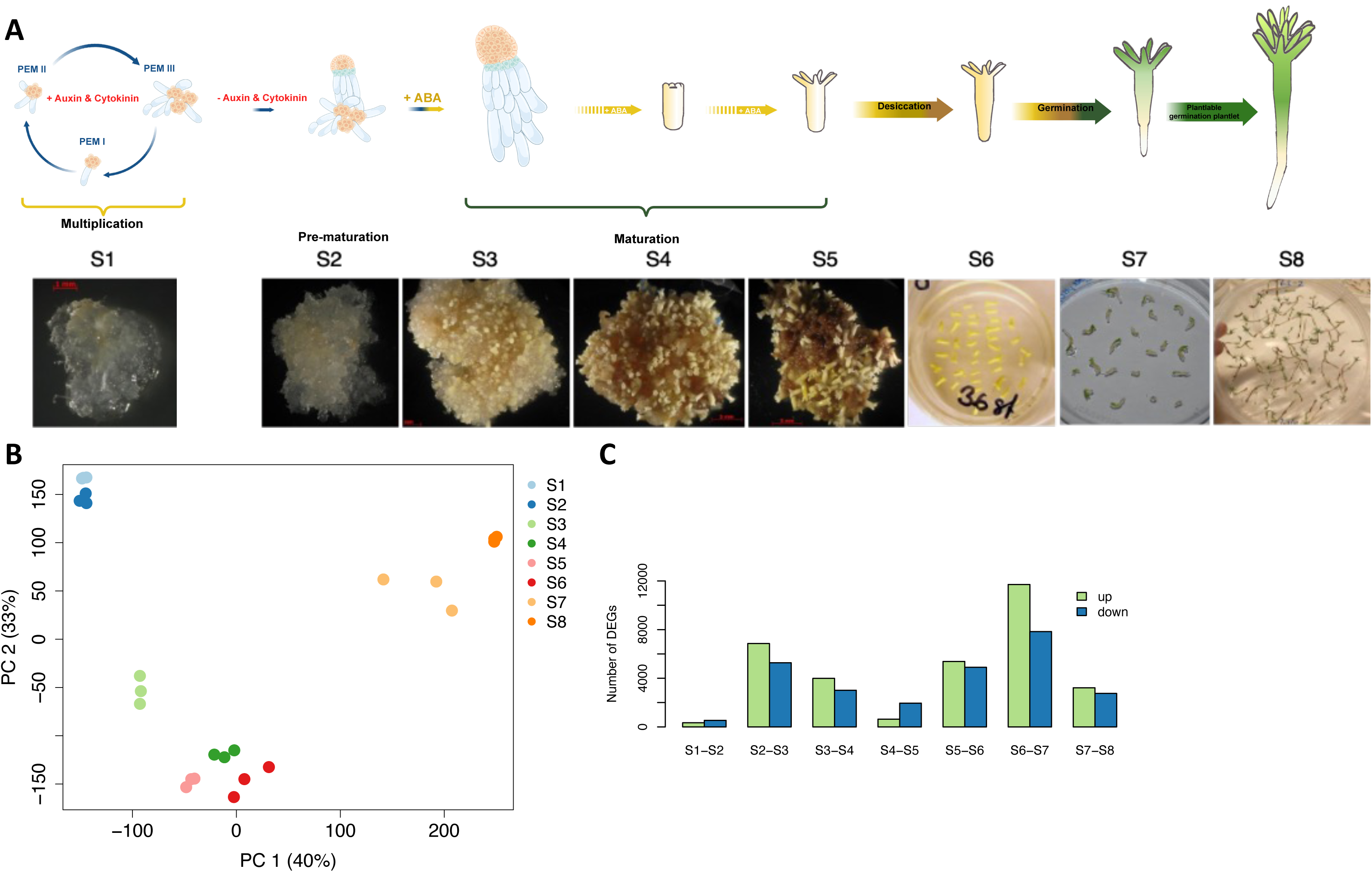
**A** Diagrammatic representation of the sampled stages during somatic embryogenesis. Representative photos of each sampled stage are shown below the diagram. Each sample consisted of a pool of hundreds of pro-embryogenic masses (PEMs; S1 and S2) that start developing into embryos (from S3). S1 corresponds to pro-embryogenic masses (PEMs) collected from proliferation medium; S2 was PEMs collected after one week on pre-maturation medium; S3, S4, and S5 were samples from the SE culture with maturing embryos collected after three, six and eight weeks on maturation medium; S6 was mature isolated embryos collected after three weeks of desiccation; S7 was mature, desiccated embryos germinated for one week to show early root formation, and S8 correspond to germinants ready to be planted and be established in compost (Fig. 1). **B** Principal component analysis plot of the RNA-Sequencing data showing the first two principal components. Colours indicate sample stage. **C** Bar graph representation of the number of differentially expressed genes (FDR-adjusted p-value <0.01 and |log2 fold change| ≥0.5). Down-regulated genes are coloured blue and up-regulated genes green.

The relationship among sampled stages was examined using PCA, in which the first two principal components (PCs) together explained 73 % of variation in the data (Figure 2B). The first PC (40 % variance explained) separated plant (S7 and S8) and embryogenic stages of the profiled development series, while the second PC (33 % variance explained) separated embryonic stages. Stages S1 and S2 clustered together, indicating high similarity in the transcriptome of samples at these two stages, which is expected as the two stages represented PEMs only distinguished by stage S2 being collected after one week on pre-maturation medium (a medium without growth regulators). Stages S3-S6, which represented embryo maturation, grouped together and formed a cluster apart from the earlier two stages. Stage S3 was the stage where the PEMs were expected to first respond to the maturation treatment, which included a higher osmoticum and ABA, by stopping multiplication and initiating maturation; Stages S4 and S5 represent a gradual maturation process until formation of a fully mature embryo. During desiccation of the mature somatic embryos, between stages S5 and S6, many processes related to dormancy in the seed are expected to take place. The third observed cluster consisted of samples from stages S7 and S8, collected after the start of germination (stage S7) and further development to plantlets that were ready to be planted into soil (stage S8). As such, the PCA revealed two major transitions in the composition of the transcriptome, the first representing a shift from proliferation to maturation (stage S2 to S3) and the second representing transition from desiccated Sem to germination (stage S6 to S7).

A total of 40281 genes were detected as expressed in SE samples of which 69.5 % (27995 genes) were DE at some point during the experiment, with the number of DEGs varying substantially between consecutive time points (Figure 2C). Three major changes in the number of DEGs were observed, with the greatest change (19544 genes) occurring during germination of desiccated embryos (S6 and S7) followed by mid stage maturing embryos compared to early maturing embryos (S2 and S3; 12,135 genes) and during desiccation (stage S5 and S6; 10282 genes). 6857 DEGs were upregulated at S2-S3 of which 1650 were downregulated and 1000 also upregulated at S5-S6. Similarly, of 5278 DEGs downregulated at S2-S3, 964 were upregulated and 737 downregulated at S5-S6. As such, it was not simply that the direction of change in expression at S2-S3 was reversed at the S5-S6 transition.

Taken together, the PCA and DE analyses revealed extensive modulation of the transcriptome during SE between stage S2 and S3 and stage S5 and S6 with a later, and expected, additional change resulting from germination and the establishment of photosynthesis. As expected, at the later transition (S6-S7) there were many significantly enriched Gene Ontology (GO) categories associated with photosynthesis (Table S2; GO:0015979 - photosynthesis (padj = 6.42e-8), GO:0005975 - carbohydrate metabolic process (padj = 1.36e-4), GO:0015994 - chlorophyll metabolic process (padj = 8.16e-3)). At the earlier transition we found groups of overrepresented genes, but the role of these was not clearly identifiable due to the general nature of the enriched GO terms (the five most enriched terms are: GO:0005975 - carbohydrate metabolic process (padj = 2.02e- 21), GO:0044260 - cellular macromolecule metabolic process (padj = 5.29e-20), GO:0045229 - external encapsulating structure organization (padj = 2.20e-18), GO:0071554 - cell wall organization or biogenesis (padj = 5.74e-12), GO:0006091 - generation of precursor metabolites and energy (padj = 8.25e-07)). This is a common challenge faced when working with non-model organisms lacking high quality functional annotations, in particular in the current case where extensive divergence from Arabidopsis makes ortholog-based annotation problematic.

### Comparison of gene expression during ZE and SE

One of the aims of the study was to identify differences in expression between SE and ZE development. There were similar numbers of genes detected as expressed in total and at each sampled stage in both datasets. As such, there was no clear indication of a general activation or transcriptome-wide mis-regulation of expression in SE compared to ZE. While the number of genes detected as expressed was similar in the two sample sets, with a relatively constant number of genes detected as expressed at all sampled stages (Figure S1), there was a clear difference in the number of DEGs, with more extensive DE between consecutive stages in SE (27995 non-redundant DEGs) than in ZE (14686 non-redundant DEGs).

Intersection of sets of genes DE between any consecutive time points in each tissue (Figure S2) revealed DEGs unique to each of the examined tissues and that Sem and Zem shared more DEGs than did FG with either type of embryo. Uniquely DEG in SE regardless of when that DE occurred were examined. These 16258 DEGs were notably enriched for genes involved in stress related responses including GO:0006952 - defence response (padj = 0.0241), GO:0050896 - response to stimuli (padj = 0.000161), GO:0055114 - oxidation−reduction (padj = 3.36E-06), GO:0016491 - oxidoreductase activity (padj = 0.000259), GO:0016209 - antioxidant activity (padj = 0.00105), GO:0072593 – reactive oxygen species metabolic processes (padj = 0.000441). The importance of stress during SE is well established (Fehér, 2015). Furthermore, DEGs reflecting the in vitro environment of exposure to plant growth regulators; GO:0008144 - drug binding (padj = 3.81E-05), GO:0009636 - response to toxic substances (padj = 0.0224), GO:0010817 - regulation of hormone level (padj = 0.0254) were also present. While these genes were not classified as DE in the ZE samples, examination of their expression suggested most were expressed in ZE with variable expression in different stages and tissues (Figure S3). The extent of overlap of DEGs at the most marked stage transitions in each sample set was examined. At the first major transcriptome transition 2611 DEGs were in common to the S2-S3 and Z3-Z4 Zem stage comparisons, representing ∼40 % of DEGs at Z3-Z4 Zem, and 753 (∼48 %) between S2-S3 and Z3-Z4 FG. Similarly, at the second major transcriptome transition, represented by S5-S6 and Z6-Z7, there were 1151 and 1071 DEGs in common between S5-S6 and Z6-Z7 Zem and Z6-Z7 FG, respectively; representing ∼50 % of DEGs in Z6-Z7 Zem and ∼30 % in Z6-Z7 FG. As such, DEGs from both ZE tissue types were DE in the SE samples despite the lack of FG tissue in SE.

There were 2949 DEGs unique to ZE regardless of when the DE occurred. These DEGs were enriched for the GO terms (Table S3) gene expression (GO:0009058, padj = 3.47e-2) and biosynthetic process (GO:0010467, padj = 3.72e-2) and were coming from all three tissue types (the early stage developing seed (SD), FG, Zem); 377 DEGs were unique to SD, 954 DEGs to Zem, 830 DEGs to FG and 788 in more than one tissue type (most often shared between the SD and Zem) (Figure 3A). DEGs unique to ZE could be important for explaining differences in SE development and could indicate which processes are crucial to achieve better performance of the SE protocol.

**Figure 3.**
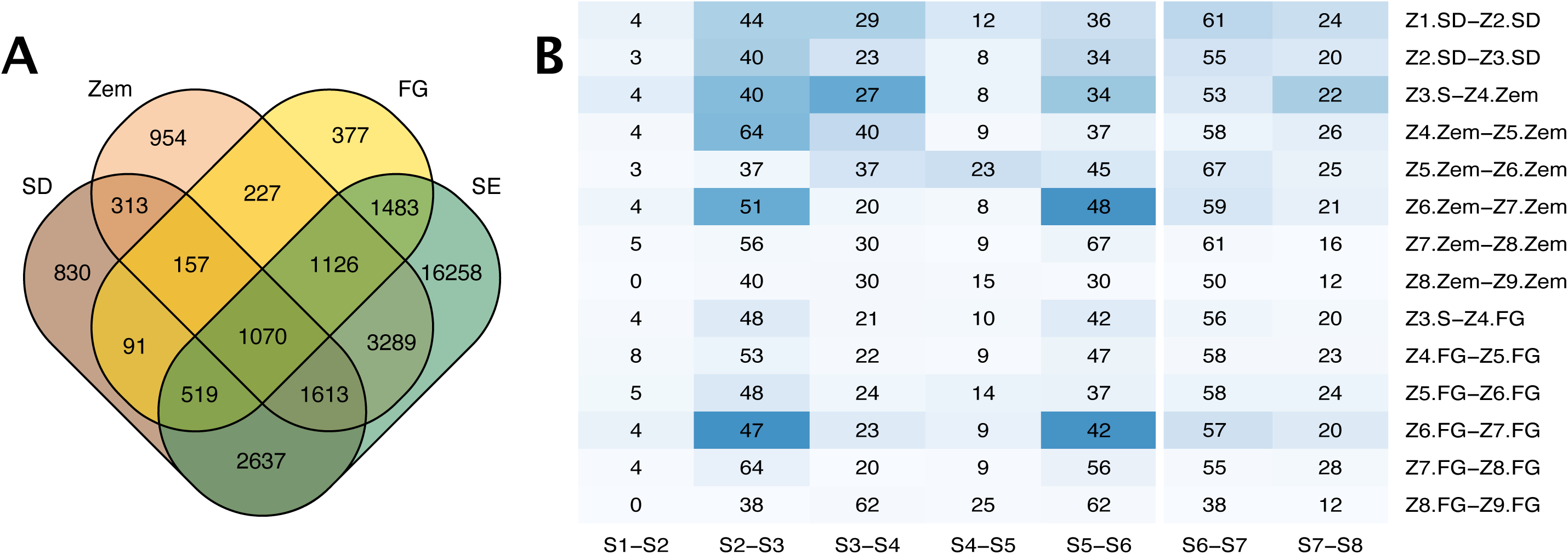
**A** Intersection of differentially expressed genes from all sample types at all stage transitions. **B** Intersection counts of differentially expressed genes at each stage transition in the zygotic and somatic embryogenesis datasets. The values indicate the percentage of differentially expressed genes detected in zygotic embryogenesis that were also differentially expressed in somatic embryogenesis. To aid visual exploration in identifying points of high similarity, cells are shaded based on -log10(p-value) from white (not significant) to dark blue (highly significant) to indicate the significance of the indicated intersection. FG, female gametophyte; Zem, zygotic embryo; SD, seed; SE, somatic embryogenesis.

Common DEGs between the transitions of SE and ZE indicated that the transition from S5 to S6, where desiccation occurred, was most similar to the transition from Z6 to Z7 in ZE (Figure 3B). In ZE, these genes were distinctly DE between Z6 to Z7 and generally remained highly expressed in subsequent sampling points. Examining the set of Zem DEGs that were also DE in SE (Figure S4A) or that were not also DE in SE (Figure S4B) revealed highly contrasting expression profiles between the two sets of samples. A number of DEGs from Zem were up-regulated across many consecutive stages but were up-regulated at only a single transition in SE (e.g., the second cluster from the top in Figure S4A). As such, there were clear contrasts in the regulation of DEGs during desiccation between the two developmental processes. Genes specifically DE in desiccated Zem and not during SE included ribosomal elements known to be associated with the storage and regulation of mRNA translation during seed dormancy (Bai et al. 2020), genes associated with mitochondrial activity related to the TCA cycle important during germination (Zhang and Fernie, 2023) and expression of genes related to the onset of germination as a morphogenic event involving cellular processes that require utilization of lipids, carbohydrates and starch. Previous analysis of quantitative data on water content, carbohydrates and total protein collected from the later stages of Zem development and Sem development in *Pinus pinaster* also indicated that the most advanced mature stage that the somatic embryos could reach in culture corresponded to the fresh, non-desiccated stage of Zem development, suggesting that the Sem maturation does not get fully completed (Morel et al. 2014). The addition of sugar alcohols at a late stage of Sem maturation has previously been shown to increase the quality of Sem, possibly by decreasing the water content in the embryo thus mimicking the desiccation process in the seed (Morel et al. 2014).

As TFs play essential regulatory roles and can have major impacts on transcriptional programmes, the sets of DE TFs in the different sample types were examined (Figure 4A). Among the set of TF families containing differentially expressed TFs in any of the samples, half were common to SD, Zem and FG (Figure 4B). Considering only ZE samples, there were more families represented in Zem than FG and genes from seven TF families were DE only in Zem samples. Families common to SD and Zem may actually be Zem specific, but it is not possible to determine which tissue(s) of the SD samples those genes were expressed in. Within the ZE dataset, a single TF family, Nuclear Transcription Factor, X-Box Binding 1 (NF-X1), was uniquely present in FG. *AtNFXL1* is component of a regulatory mechanism that has been shown to improve physiological status and to support growth and survival under stress (Müssig et al. 2010). All TF families DE in the ZE samples (of all types) were DE in SE in addition to two families (S1Fa-like and Hordeum Repressor Transcription HRT-like) that were uniquely DE in SE. Overexpression of *S1Fa* genes has been linked to improved drought tolerance in *Populus trichocarpa* (Zhao et al. 2021). Relatively little is known about *HRT-like* genes but work in barley indicated that HRT can repress the ɑ-amylase promoters (Raventós et al. 1998) and analysis of promoter binding motifs in *Erograstis tef* indicated that HRT may be involved in the control of hormones (ABA, Gibberellic acid), sugars (zein, ɑ-amylase) and abiotic stress (Mulat and Sinha, 2020). In general, the representation of TF families at different stage transitions was far more even in SE than in ZE and there were more DE TFs at more stage transitions in SE than in ZE. In the overall comparison across development, the pattern of expression for different TF families appeared more uniformly across SE than ZE with almost all families showing DE at all SE developmental transitions. In contrast, in ZE some transitions had only one (Z4.FG−Z5.FG and Z7.FG−Z8.FG), two (Z8.Zem−Z9.Zem) or three (Z7.Zem−Z8.Zem) TF families represented. This reflected the low numbers of DEGs at these stages (Figure 4A). It is also notable that in ZE, the greatest number of DEGs was identified at the transitions from polyembryogeny to early embryogeny (between Z1-Z2, Z2-Z3, and Z3-Z4, respectively), and later early to late embryogeny (Z6-Z7), which was also reflected in the extent of DE of TFs. In SE, there was no corresponding peak in DE at the transition from polyembryogeny to early embryogeny (Figure 1C and Figure 2C), indicating that prior developmental processes required for subsequent processes may not be correctly activated by the culture conditions present at the earlier stage. The importance of abiotic factors early in embryo development on the later stages of development was also recently shown for *Abies nordmanniana* (Nielsen et al. 2022), suggesting that efforts to optimize SE culture conditions should consider the interrelation between developmental and culture conditions at steps of the SE protocol. While there were no TF families uniquely DE in ZE, there were 12 TF genes that were uniquely DE in Zem (Figure 4C). While these 12 genes were not classified as significantly DE in SE they did have clear evidence of expression level variation during SE within specific stages, indicative of expression regulation. The expression profiles of these genes were distinctly different during SE than ZE and, as such, these represent interesting candidate TFs that could explain differences in the regulation of gene expression during the two developmental processes. Taken together, this suggests that the regulation of many TFs differs during the two developmental processes.

**Figure 4.**
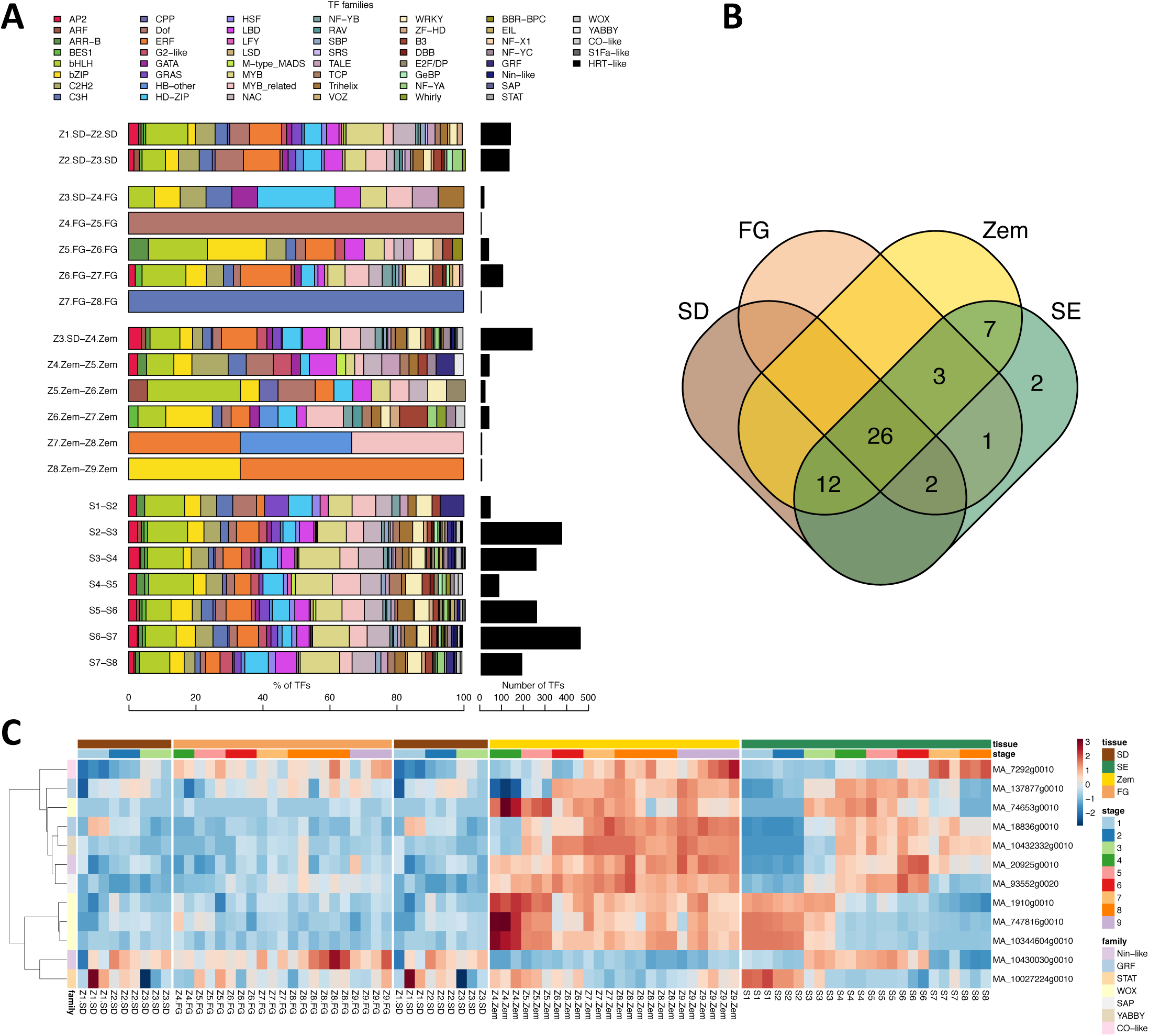
Differential expression of transcription factor genes during development of seed tissue, zygotic embryos, female gametophytes and somatic embryos. **A** The percentage representation of differentially expressed transcription factor genes in transcription factor families (left panel, coloured bars coloured according to family as indicated in the key above the bars) and corresponding total number of differentially expressed transcription factor genes (right panel, black bars) at all sample transitions. **B** Intersection of transcription factor families with differentially expressed genes in any of the sampled stages from all sample types. **C** A heatmap representation of the 12 transcription factor genes that were uniquely differentially expressed in zygotic embryo. Coloured bars above the heatmap represent sample type and sample stage, and to the left transcription factor family, as indicated in the colour legend to the right of the heatmap. The data is row-scaled (z-scores). FG, female gametophyte; Zem, zygotic embryo; SD, seed; SE, somatic embryogenesis.

### Genes of known importance for embryogenesis

A set of genes that have previously been identified as important during the process of embryogenesis was identified from the literature (Table S4) and their expression patterns (Figure S5) and differential expression between developmental stages (Table S4) were examined within the current ZE and SE data. In addition, specific TFs of known importance during embryo development and with contrasting expression between the two developmental processes were further considered.

The only TF family active in FG at the transition Z4.FG−Z5.FG represented the TCP (*Zea mays* TEOSINTE BRANCHED1 (TB1), *Antirrhinum majus* CYCLOIDEA (CYC), *Oryza sativa* PROLIFERATING CELL NUCLEAR ANTIGEN FACTOR1/2) TF family that have been implicated as regulators during plant growth, development and abiotic stress (Nie et al. 2022). TCP TFs affect growth directly via the cell cycle and indirectly via influencing plant hormonal signalling and the circadian clock (Danisman, 2016). In Arabidopsis AtTCP14 has been shown to specifically regulate embryonic growth development during germination. It is expressed primarily in the embryo with limited expression in the endosperm (Tatematsu et al. 2008). This contrasts to the observed expression pattern in the current data, where the highest sequence similarity homolog was the only DE TF at the Z4.FG−Z5.FG transition. Forty-three *TCP* genes were identified from *Pinus tabuliformis* with several being co-expressed with clock genes and during seasonal oscillations (Nie et al. 2022). These observed expression patterns could reflect function in FG during dormancy and before onset of germination.

The Ethylene Response Factor (*ERF*) family was one of two (Z8.Zem−Z9.Zem) or three (Z7.Zem−Z8.Zem) TF families that increasingly dominated the DE set of TFs towards the end of the Zem developmental process (Fig. 4A). Functional enrichment results suggested that AP2/ERFs were transcribed in the FG. There was generally little DE of TFs at these stages, which is perhaps not surprising given that the seeds were dormant. Desiccation is an essential part of the later stages of seed development, preparing for dormancy until germination is triggered. Ethylene is known to release dormancy in a number of plants (Corbineau et al. 2014) and several ERFs also bind to dehydration-responsive elements (Müller and Munné-Bosch, 2015). During the corresponding stages of SE, where stage S6 represented somatic embryos desiccated for three weeks and S7 embryos after one week in germination, there was no substantial representation of DE *ERFs*. This may be interpreted as an indication of another step of the SE process that deviates from Zem development and suggests further studies are warranted into expanding culture condition optimizations to include the gas phase of the culture containers during SE germination. Maturation success rates of Sem have been associated with production of ethylene in a number of conifer species, e.g., *Pinus sylvestris* (Lu et al. 2011), *Araucaria angustifolia* (Jo et al. 2014), *Picea glauca* (Meskaoui et al. 2000) and *Picea mariana* (Meskaoui and Tremblay, 2001). However, further studies into the effect of ethylene during germination are currently lacking. Overall, the less distinct regulation of the *ERF* family during SE may reflect missing effects of ethylene signalling during development due to Sem development occurring inside a culture container surrounded by a relatively large volume of air compared to Zem development, which is contained within the seed coat.

It is notable that the WUSCHEL -related homeobox (*WOX*) family, being the best studied TF family in conifer embryogenesis based on its function during the formation of the embryogenic shoot meristem in Arabidopsis and other plants (Hassani et al. 2022), was scarcely represented in the set of TF families containing DE genes in both ZE and SE samples (Figure 4A). However, when analysing the expression patterns of the twelve TF genes that were DE specifically in Zem and in no other tissue, the most significant changes in expression occurred for three *WOX* genes expressed in Zem during the transition from early to late embryogeny. In Sem, the expression was not enhanced during the corresponding stages of S3 and S4 (pre-maturation and early maturation) (Figure 4C). In *Pinus pinaste*r, *WOX2* was mostly expressed during proliferation of Sem (Hassani et al. 2022) and low during maturation and germination which also agrees with the three *WOX* genes in the present study.

A notable discrepancy in the expression pattern of the DE TFs in ZE and SE was observed for three putative baby boom (*BBM*) genes that in other plant species have been demonstrated to have a significant effect on induction of SE (MA_86195g0010, MA_98095g0010, MA_121578g0010 in Figure S5 and Table S4). In *Larix decidua*, a conifer species typically showing high success rates for SE plant formation, expression of *LdBBM* increased from proliferation to fully mature embryos both during ZE and SE (Rupps et al. 2016). In the present study, Zem showed a slight increase in expression over maturation similar to the observed patterns in *Larix* Zem. In contrast, expression in Sem did not follow this pattern (Figure S5), as was similarly observed in *Larix* Sem.

The signal transduction pathway involved in controlling ABA levels during embryo development involve *Viviparous-1* (*VP1*) and *VP2* (Kermode, 2005). The expression of *VP1* and *VP2* increased from early embryogeny until mid-late embryogeny similarly during Zem and Sem development (S4; MA_66505g0010 and MA_66505g0020). This is in agreement with previous findings for both ZE where the endogenous levels of ABA during Zem development in *Picea glauca* were shown to increase from relatively low levels at the onset of embryo development to peak mid stage embryo maturation (Kong et al. 1997) and the expression levels of *VP1* during *P. abies* SE (Uddenberg et al. 2011).

SE is increasingly utilized for propagation of commercially important conifers; mostly spruces in the northern hemisphere (*Picea abies*, Picea *glauca*) and pine in the southern (*Pinus radiata*). The method has potential for cost-effective, large-scale propagation through automation of mainly the *in vitro* steps (*ex vitro* steps are already automated), however the process from initiation of PEMs to the plant growing in the nursery is still far from being as efficient as the zygotic seed developmental process. To improve the survival rates and quality of embryos through development, efforts have been made to understand the natural conditions for development of the Zem by analysing the composition of the FG and then to duplicate the seed nutritional, hormonal and environmental conditions *in vitro* (reviewed by Pullman and Buccalo 2014). The data presented here cover more developmental stages than any previous study, providing a powerful resource for elucidating new knowledge of gene expression events during the very first stages of seed development, key events such as desiccation, and the differences between ZE and SE. Exploration of the data suggests more extensive differences in DEGs between the SE process and FG than between Zem and FG, with these contrasting examples of developmental regulation representing candidate genes that may link to targets for protocol improvement such as modifications to culture media and controlling gas composition during the gas phase.

## Conclusion

The comparison of ZE and SE presented here highlights that while there is extensive commonality in transcriptional regulation during ZE and SE, there are also numerous differences in the sets of genes classified as significantly differentially regulated at stage transitions during the two developmental processes in addition to differences in the timing and duration of expression changes. Genes with contrasting regulation include many TFs, which are important targets for understanding the regulatory mechanisms underlying differences in ZE and SE. Importantly, genes with contrasting regulation between ZE and SE can help identify associated biochemical, hormonal, and physiological influences and these can direct future optimization of SE protocols. For example, there were obvious differences in the timing and duration of changes in gene expression associated with desiccation. Future exploration of the known factors determining regulation of these genes could identify specific targets for optimization such as changes in the timing of application or composition of hormones in culture media or changes in environmental conditions to induce physiological responses. Such differences can help explain biological differences in the *in vitro* and *in planta* processes, which helps provide greater understanding of which regulatory events are important in the natural process of Zem in conifers. Exploration of this data resource identified essential factors in the natural seed maturation process, as summarized by the following findings: 1) further evidence to support the hypothesis that the desiccation process in Sem is incomplete compared to this process during maturation in Zem, thereby providing details on key regulatory pathways required for ZE development. These findings point to the importance of the interrelation of abiotic conditions during embryo development for the completion of the developmental process and provide information useful for directing optimizations of SE protocols: 2) the expression of BBM may be used as an indicator of successful Sem and, as such, may represent a key regulatory mechanism in embryogenesis; 3) specific transcription factors with contrasting expression were identified between Zem and Sem development, which represent candidates for future characterisation studies to more fully determine their role in regulating conifer embryogenesis. This data resource, which is explorable within expression visualisation tools at the PlantGenIE.org resource, provides a comprehensive reference resource for the plant community for advancing understanding of the process of SE, including for comparative studies between gymnosperms and angiosperms, and for identifying the regulatory basis of differences between SE and ZE.

## Experimental procedures

### Plant material

Material for the ZE samples was obtained from cones collected from one Norway spruce (*Picea abies* (L.) Karst) tree that was control pollinated with a known mixed pollen source at the Swedish Forest Research Institute (Skogforsk) in Sävar, Sweden, starting from two weeks after pollination on June 4^th^, 2017, then every two weeks until full cone maturity on September 27^th^, 2017. The cones were stored at +4°C until sample extraction within a week from collection.

Embryogenic cultures were initiated and captured in 2014 from seeds from the Köttsjö Norway spruce tree, Z4006 (Nystedt et al. 2013). Cultures from cell line K14-03 were proliferated for embryo maturation and germination by standard methods (Arnold and Clapham 2008). Briefly, proliferation was stimulated using liquid half-strength LP medium supplemented with auxin (2,4-D; 2.21 mg l−1), N6-benzyladenine (BA; 1 mg l−1) and sucrose (10 g l−1). After 4–6 weeks cultures were transferred to liquid pre maturation medium devoid of 2,4-D and BA for one week, then to liquid maturation medium supplemented with ABA (16 mg l^−1^) and sucrose (30 g l^−1^) for six weeks. Cultures were kept in darkness at 20 °C. Mature embryos were dried for three weeks under high humidity in darkness and subsequently germinated in 15 cm Petri dishes on solid medium containing minerals and vitamins supplemented with sucrose (30 g l^−1^) and casein hydrolysate (0.5 g l^−1^), solidified with 3.5% Gelrite. The pH was adjusted to 5.8 ± 0.1 prior to autoclaving. Germinants were grown for three weeks in continuous red light (wavelength: 660 nm; TL-D 18W/15, Philips, Stockholm, Sweden) at 5 μmol m−2 s^−1^ at 20 °C, then moved under continuous white fluorescent tubes (Fluora L 18W/77, Osram, Johanneshov, Sweden) at 100–150 μmol m^−2^ s^−1^ at 20 °C (Kvaalen and Appelgren 1999). Germinants with actively growing shoot and root were planted in peat moss, acclimatized and plant growth established under the same light conditions.

### Sampling

From the three earliest cone-collection times, the whole female gametophyte (FG) with the seed coat removed was sampled as no zygotic embryo (Zem) could be isolated. The two earliest collection times (Z1-2) had distinctly smaller FG compared to collection three (Z3). From the fourth collection time on July 31^st^ (Z4) until ninth collection time (Z9), samples were separated into zygotic embryos (Z4.Zem) and FG (Z4.FG). A total of 1618 seeds were collected during the duration of the sampling period whereof 696 whole FGs were collected from Z1-3 and 922 FMG and 838 Zem from stages Z3-Z10. A minimum of 100 samples were collected from each collection date. The samples were mixed and split into separate Eppendorf tubes with approximately 25 samples of zygotic embryos with FG for stages Z1-2, and subsequently zygotic embryos or FGs, snap-frozen in liquid nitrogen and stored at -80 °C until further processed.

Due to insect damage of cones, limited numbers of samples were collected on Oct 11th (Zem9/10 and FG9/10). In addition to the sample collection from the controlled pollinated tree, dried seeds collected in 2016 from a related tree were sampled and used as additional Zem and FG replicates at stages 9 and 10 (see associated metadata at ENA).

Samples were collected from eight different stages of somatic embryo development from pro-embryogenic masses to SE germinants established in compost as follows: S1 corresponds to pro-embryogenic masses (PEMs) collected from proliferation medium; S2 was PEMs collected after one week on pre-maturation medium; S3, S4, and S5 were samples from the SE culture with maturing embryos collected after three, six and eight weeks on maturation medium; S6 was mature isolated embryos collected after three weeks of desiccation; S7 was mature, desiccated embryos germinated for one week to show early root formation, and S8 correspond to germinants ready to be planted and be established in compost (Fig. 1). Samples were collected in 2014. For each sampling point, 50 mg of somatic embryo tissue (PEMs, mature embryos, germinants, or plants) were collected from each of three different parallel petri plates for every developmental stage. Samples from each developmental stage were mixed and split into four Eppendorf tubes, snap-frozen in liquid nitrogen and stored at -80°C until further processed.

### RNA isolation and sequencing

Total RNA was extracted from the ground samples with an RNeasy Mini Kit (Qiagen, Hilden, Germany) according to the manufacturer’s guidelines. Nucleotide quantity and purity were assessed using a NanoDrop 2000C Spectrophotometer (Thermo Scientific, Waltham, MA, USA) and quality of total RNA using an Agilent 2100 Bioanalyzer (Agilent Technologies, Inc., Santa Clara, CA, USA). RNA libraries for sequencing were prepared using TruSeq Stranded mRNA sample prep kit with 96 dual indexes (Illumina, San Diego, CA, USA) according to the manufacturer’s instructions with the following changes: RNA was fragmented 5’ and the protocols were automated using an Agilent NGS workstation (Agilent Technologies) with purification steps described by Lundin et al. (2010) and Borgström et al. (2011).

### Transcriptome sequencing and data pre-processing

RNA isolated from Zem and FG collected at eight time points during seed development (with the ZE and FG processed together for the first three collection times) and eight developmental stages during somatic embryogenesis (SE) was sequenced on Illumina HiSeq 2500 at SciLifeLab using 2x126 bp paired-end reads to an average read number of 24.1 ± 2.1 M reads per sample. Three biological replicates were sequenced for each time point/developmental stage.

The quality of the raw sequence data was assessed using FastQC (http://www.bioinformatics.babraham.ac.uk/projects/fastqc/), v0.11.4. Sequence reads originating from ribosomal RNAs (rRNA) were identified and removed using SortMeRNA (v2.1; Kopylova et al. 2012; settings --log --paired_in --fastx--sam --num_alignments 1) using the rRNA sequences provided with SortMeRNA (rfam-5s-database-id98.fasta, rfam-5.8s-database-id98.fasta, silva-arc-16s-database-id95.fasta, silva-bac-16s-database-id85.fasta, silva-euk-18s-database-id95.fasta, silva-arc-23s-database-id98.fasta, silva-bac-23s-database-id98.fasta and silva-euk-28s-database-id98.fasta). Data were then filtered to remove adapters and trimmed for quality using Trimmomatic (v0.36; Bolger *et al*. 2014; settings TruSeq3-PE-2.fa:2:30:10 SLIDINGWINDOW:5:20 MINLEN:50). After both filtering steps, FastQC was run again to ensure that no technical artifacts were introduced. Read counts were obtained using Salmon (v. 0.11.2; Patro et al. 2017) using the *P. abies* v1.0 transcript sequences as a reference (retrieved from the PlantGenIE resource; Sundell *et al*. 2015).

### Differential expression analysis of genes

The salmon abundance values were imported into R (v4.0.0; R Core Team 2015) using the Bioconductor (v3.11; Gentleman et al. 2004) tximport package (v.1.16.1; Soneson et al. 2015). For the data quality assessment (QA) and visualisation, the read counts were normalised using a variance stabilising transformation as implemented in the Bioconductor DESeq2 package (v1.28.1; Love et al. 2014). The biological relevance of the data - e.g. biological replicates similarity - was assessed by Principal Component Analysis (PCA) and other visualisations (e.g. heatmaps), using custom R scripts, which identified one sample as an outlier, which was therefore removed from further analysis. Statistical analysis of gene differential expression (DE) between consecutive stages of the experiment was performed in R using the Bioconductor DESeq2 package. FDR adjusted p-values (by the Benjamini–Hochberg procedure) were used to assess significance; a common adjusted threshold of 1% was used throughout. All expression results were generated in R, using custom scripts. Formulas used in DESeq2 included stages and DEGs between consecutive stages of the experiment were extracted using the function ‘results’, provided with option ‘filter = rowMedians(counts(dds)’. DEGs considered for further analysis were filtered by fold change |log2FC| ≥ 0.5 and p-value adjusted for multiple testings padj < 0.01. The number of genes in the intercept whose padj value was different from NA was considered to be the number of all the expressed genes in the experiment.

### Functional enrichment analysis

Gene Ontology (Ashburner *et al*. 2000) functional enrichment of differentially expressed genes (DEGs) and clusters of co-expressed genes at *P* < 0.05 was analysed using an in-house implementation of the parent-child test (Grossmann *et al*. 2007) and a Benjamini-Hochberg multiple testing correction. The background set of genes used for the test were all the genes expressed in the experiment or union of the genes expressed in SE and ZE when comparing groups of DEGs from different experiments.

### Data statement

Raw sequencing data is available from the European Nucleotide Archive (ENA) as accession PRJEB72619.

R scripts used to perform bioinformatic analysis are available in the GitHub repository (https://github.com/Emkago/spruce-embryogenesis-mRNA).

Gene expression data have been integrated at PlantGenIE.org into the available expression visualisation tools. All results of differential expression test, the normalised gene expression values used to perform the presented analyses and all functional enrichment test results are available at the Science for Life (SciLife) data centre FigShare resource doi:10.17044/scilifelab.25315867.

## Accession numbers

PRJEB72619

## Acknowledgements

Iftikhar Ahmad for collecting samples for the zygotic embryogenesis series and Ioana Gaboreanu and Sofie Johansson for collecting samples for the somatic embryogenesis series. Katja Stojkovič was supported by a grant from the Kempe Foundation (SMK1340). Nathaniel Street and Ulrika Egertsdotter are supported by the Trees for the Future (T4F) project. This work was supported by grants from the Knut and Alice Wallenberg Foundation. We thank the UPSC Bioinformatics facility for support. The authors acknowledge support from the National Genomics Infrastructure in Genomics Production Stockholm funded by Science for Life Laboratory, the Knut and Alice Wallenberg Foundation and the Swedish Research Council, SNIC/Uppsala Multidisciplinary Center for Advanced Computational Science for assistance with massively parallel sequencing and access to the UPPMAX computational infrastructure, and the Umeå Plant Science Centre bioinformatics facility for support.

## Short legends for Supporting Information

**Figure S1** Number of genes with any aligned sequencing read and classified as expressed at different expression thresholds (increasing variance stabilising transformation values) in the samples of zygotic embryogenesis (**A**) and somatic embryogenesis (**B**)

**Figure S2** Intersection of genes differentially expressed between all consecutive stages from each tissue analysed during somatic and zygotic embryogenesis. FMG, female gametophyte.

**Figure S3** Expression of genes in zygotic embryogenesis samples that were uniquely differentially expressed during somatic embryogenesis. Expression values for each gene were Z-score normalized across samples. FG, female gametophyte; Zem, zygotic embryo; SD, seed; Z1-Z9 sampling stages from zygotic embryogenesis. Each sample from Z1 – Z3 is represented twice for technical reasons.

**Figure S4 A** Expression of genes differentially expressed in both somatic and zygotic embryogenesis during desiccation of the zygotic embryo (stage Z6 to Z7). **B** Expression of genes differentially expressed at transition Z6-Z7 in zygotic embryos but not differentially expressed in somatic embryogenesis. Expression values for each gene were Z-score normalized across samples. FG, female gametophyte; Zem, zygotic embryo; SD, seed; SE, Somatic embryogenesis; Z1-Z9, sampled stages during zygotic embryogenesis; S1-S8, sampled stages during somatic embryogenesis.

**Figure S5** Expression of genes known to be important in embryogenesis in the processes of somatic embryo initiation, desiccation and nitrogen utilisation shown in somatic and zygotic embryogenesis samples. Expression values for each gene were Z-score normalized across samples. FG, female gametophyte; Zem, zygotic embryo; SD, seed; SE, Somatic embryogenesis; Z1-Z9, sampled stages during zygotic embryogenesis; S1-S8, sampled stages during somatic embryogenesis. Details is the genes are available in Table S4.

**Table S1** Functional enrichment of differentially expressed genes during zygotic embryogenesis.

**Table S2** Functional enrichment of differentially expressed genes during somatic embryogenesis.

**Table S3** Functional enrichment of differentially expressed genes between somatic and zygotic embryogenesis or for different zygotic tissues.

**Table S4** Differential expression in zygotic and somatic embryogenesis stage transitions of genes known to be important in embryogenesis.

