## Supplementary figures and images for "A transcriptome atlas of zygotic and somatic embryogenesis in Norway spruce"

### Figure S1

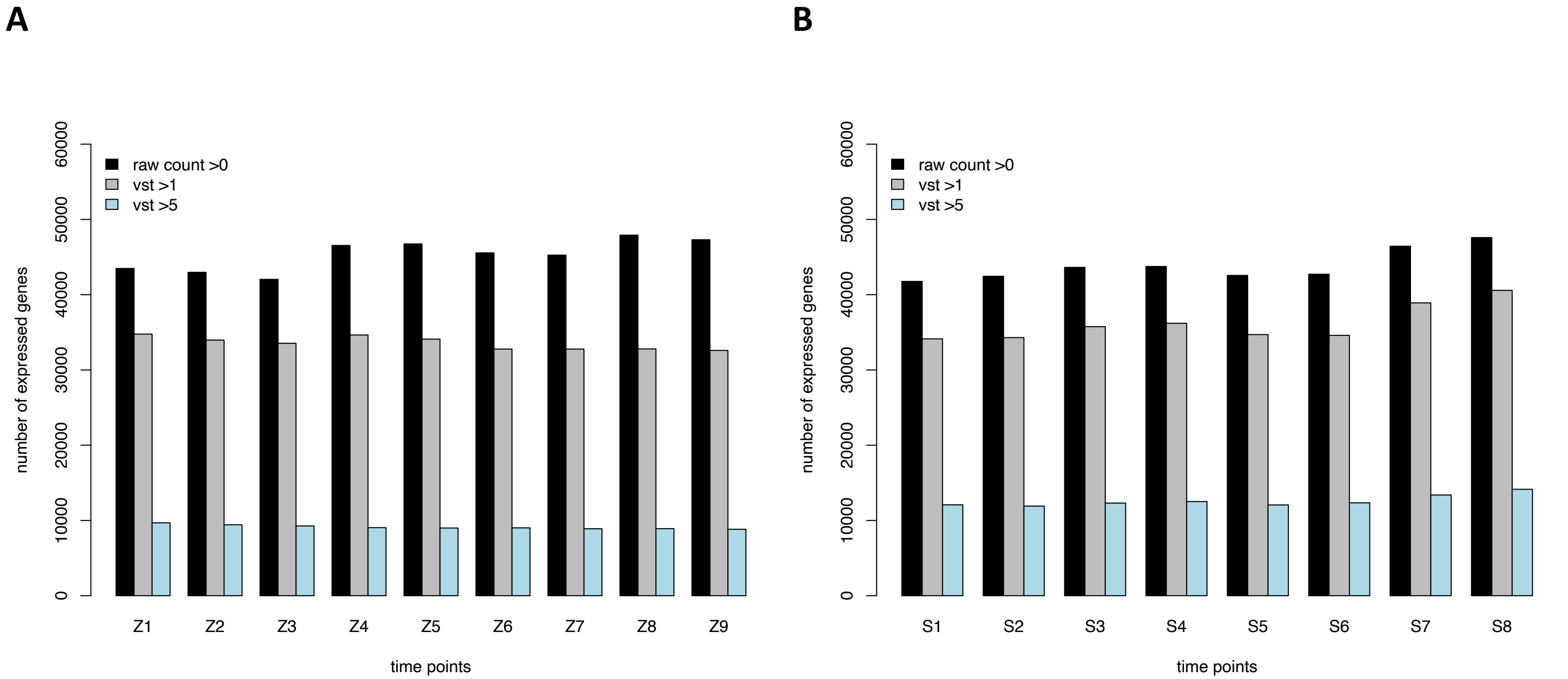

### Figure S2

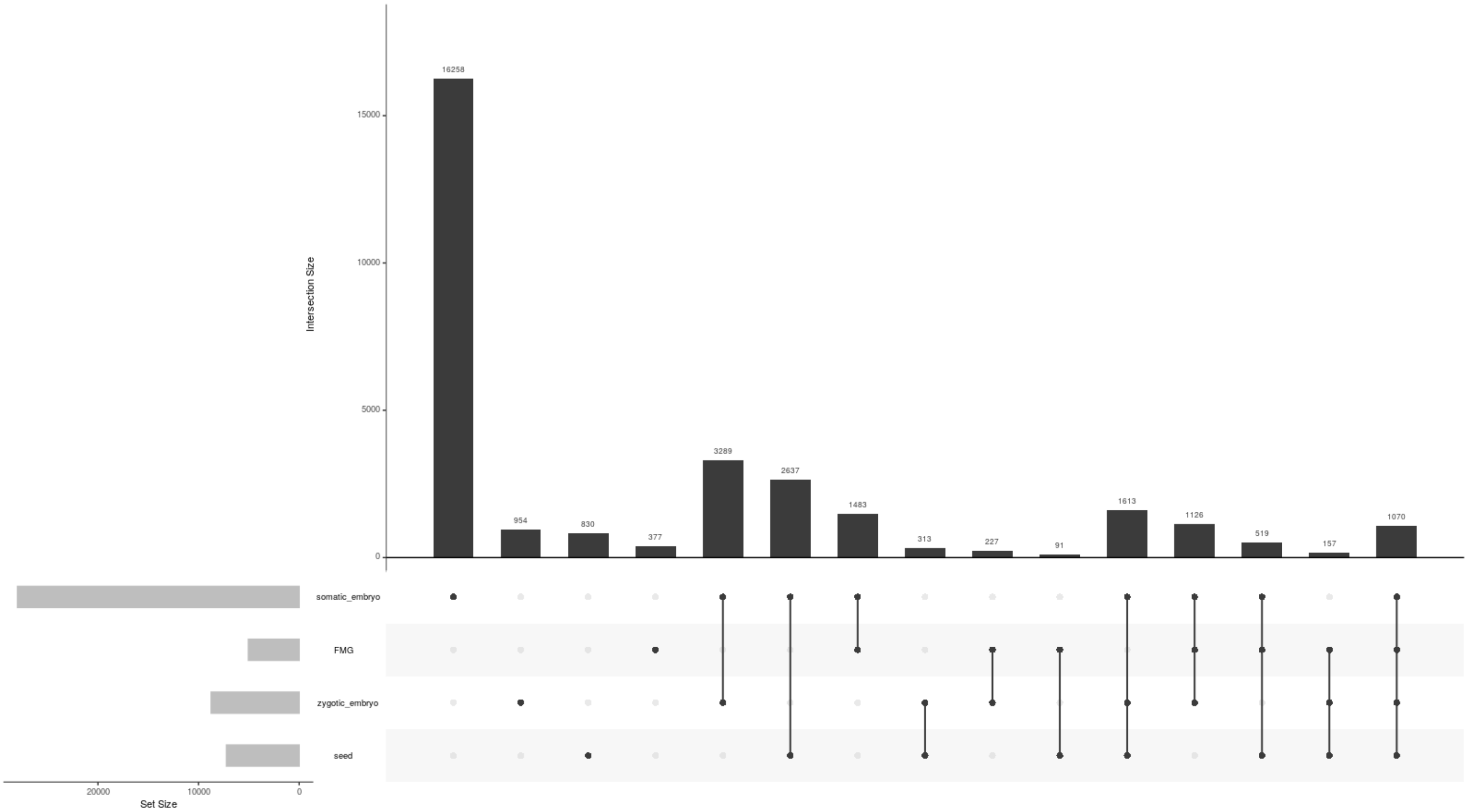

### Figure S3

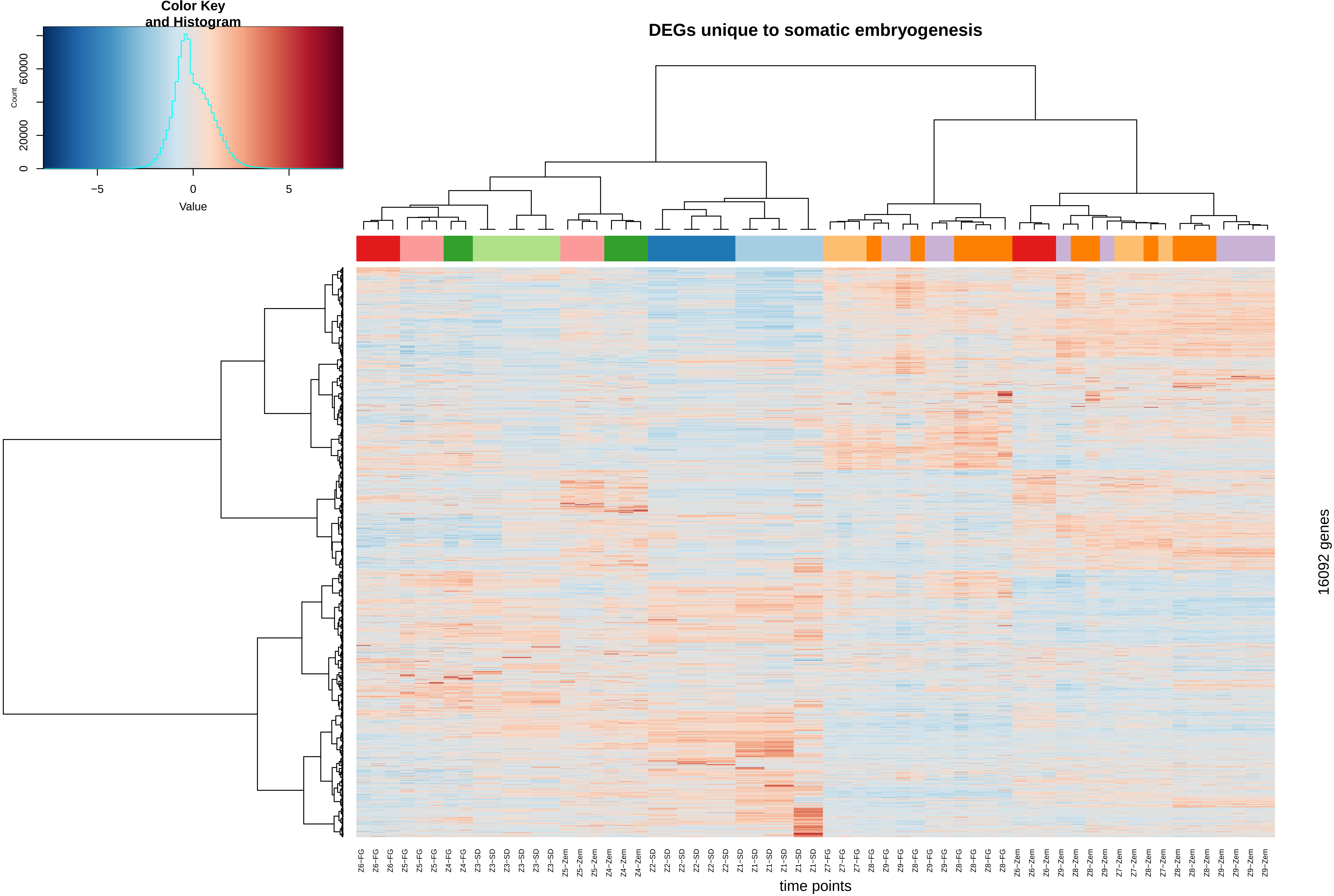

### Figure S4

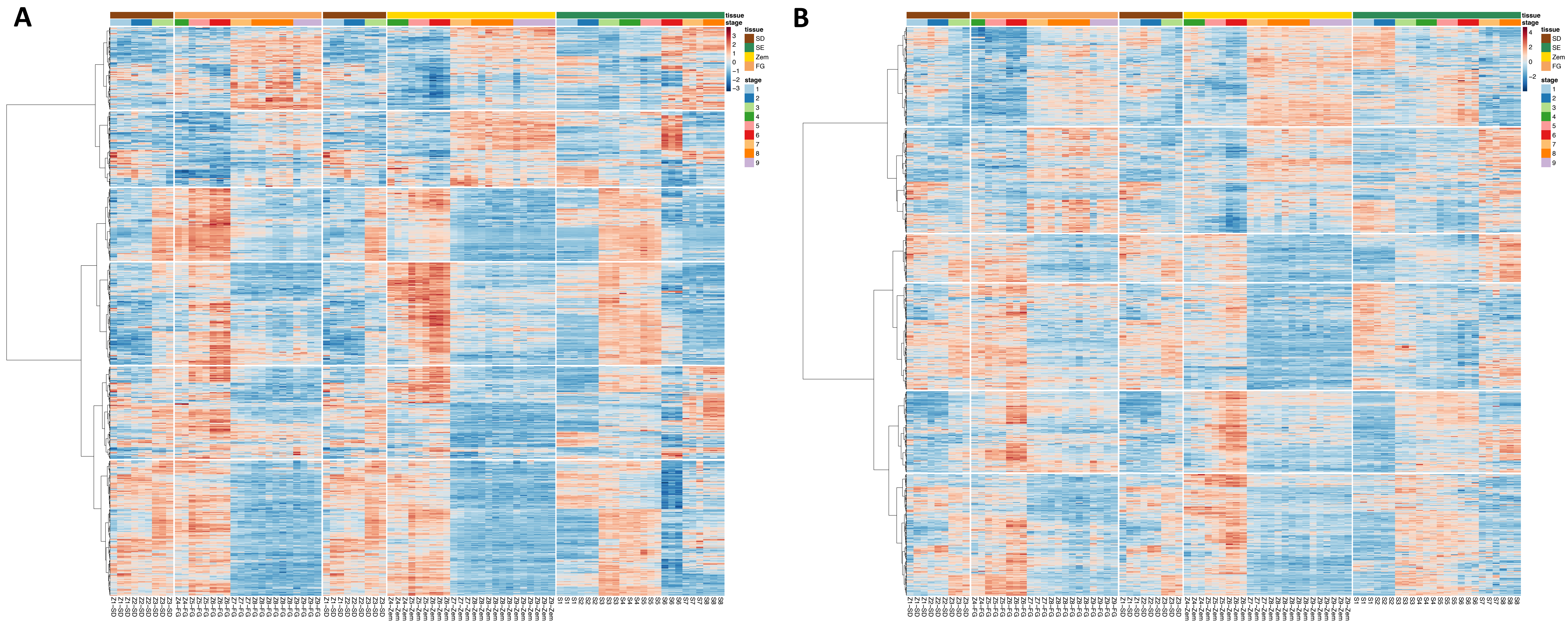

### Figure S5

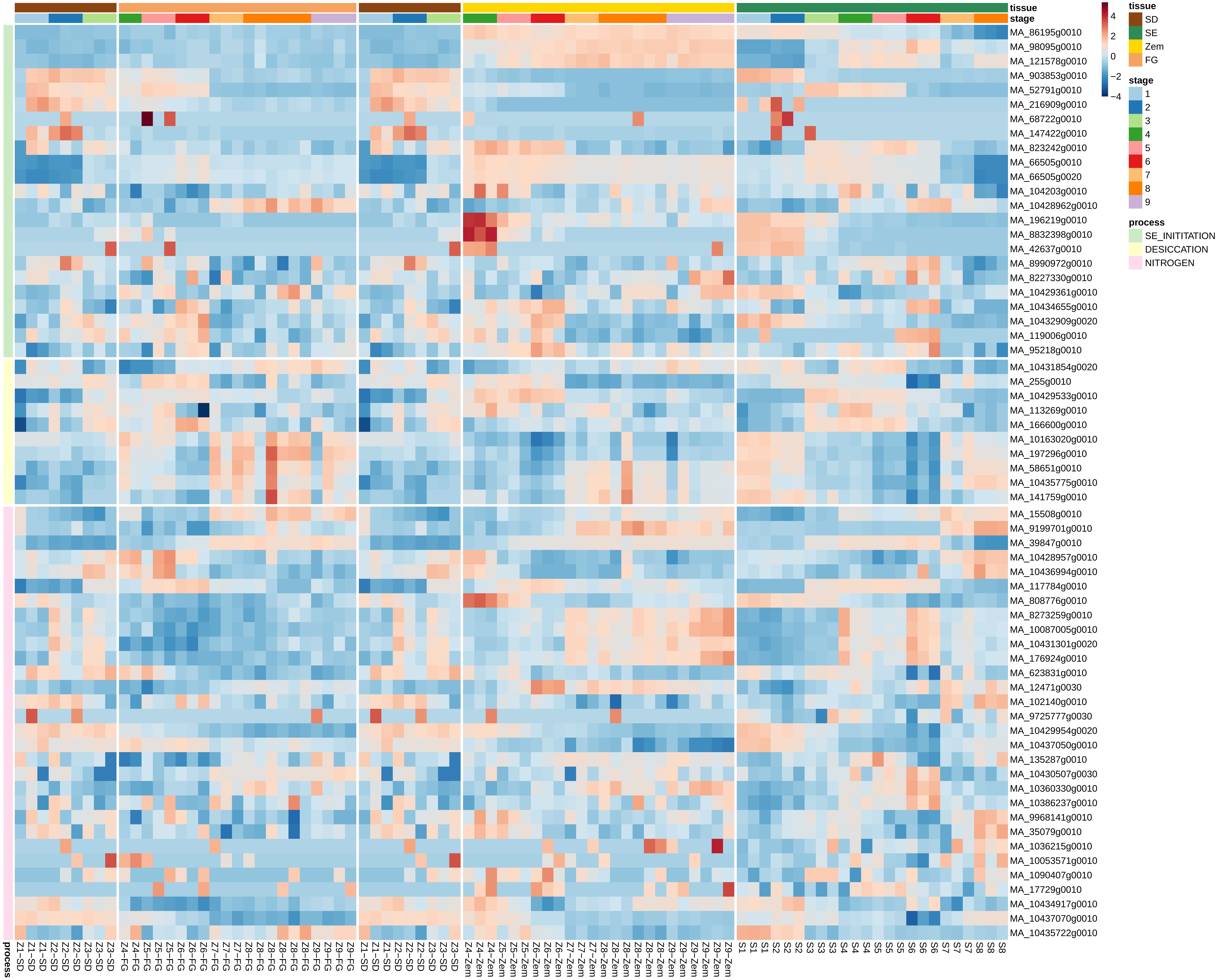
